# Conformational flexibility is a critical factor in designing broad-spectrum human norovirus protease inhibitors

**DOI:** 10.1101/2024.09.16.613336

**Authors:** Son Pham, Boyang Zhao, Neetu Neetu, Banumathi Sankaran, Ketki Patil, Sasirekha Ramani, Yongcheng Song, Mary K. Estes, Timothy Palzkill, B.V. Venkataram Prasad

## Abstract

Human norovirus (HuNoV) infection is a global health and economic burden. Currently, there are no licensed HuNoV vaccines or antiviral drugs available. The protease encoded by the HuNoV genome plays a critical role in virus replication by cleaving the polyprotein and is, therefore, an excellent target for developing small molecule inhibitors. While rupintrivir, a potent small-molecule inhibitor of several picornavirus proteases, effectively inhibits GI.1 protease, it is an order of magnitude less effective against GII protease. Other GI.1 protease inhibitors also tend to be less effective against GII proteases. To understand the structural basis for the potency difference, we determined the crystal structures of proteases of GI.1, pandemic GII.4 (Houston and Sydney), and GII.3 in complex with rupintrivir. These structures show that the open substrate pocket in GI protease binds rupintrivir without requiring significant conformational changes, whereas, in GII proteases, the closed pocket flexibly extends, reorienting arginine-112 in the BII-CII loop to accommodate rupintrivir. Structures of R112A protease mutants with rupintrivir, coupled with enzymatic and inhibition studies, suggest R112 is involved in displacing both substrate and ligands from the active site, implying a role in the release of cleaved products during polyprotein processing. Thus, the primary determinant for differential inhibitor potency between the GI and GII proteases is the increased flexibility in the BII-CII loop of the GII proteases caused by H-G mutation in this loop. Therefore, the inherent flexibility of the BII-CII loop in GII proteases is a critical factor to consider when developing broad-spectrum inhibitors for HuNoV proteases.

**IMPORTANCE:** Human noroviruses are a significant cause of sporadic and epidemic gastroenteritis worldwide. There are no vaccines or antiviral drugs currently available to treat infections. Our work elucidates the structural differences between GI.1 and GII proteases in response to inhibitor binding and will inform the future development of broad-spectrum norovirus protease inhibitors.

## INTRODUCTION

Acute gastroenteritis is a significant global health issue, particularly affecting infants and young children in low and middle-income countries. After rotavirus vaccines became widely used, norovirus emerged as the most common cause of viral gastroenteritis^1,2^. Norovirus also severely affects immunocompromised and older adults, who account for the majority of mortalities in high-income countries^3^. Globally, there are over 685 million cases of norovirus infection annually, resulting in an estimated $64 billion in combined medical and societal costs each year^4^.

Noroviruses are divided into ten genogroups (GI to GX), and each genogroup further into several genotypes^5,6,7^. More than 30 genotypes among GI, GII, GIV, GVIII, and GIX genogroups infect humans^8^. Among these human noroviruses (HuNoVs), the GII.4 genotype (genogroup II and genotype 4) causes the most infections globally, with GII.2 and GII.17 strains causing localized epidemics^9,10^. The GII.4 variants have been responsible for 80% of global outbreaks, including six pandemics^11^. The GII.4 HuNoVs undergo epochal evolution with new variants with different antigenic profiles emerging periodically^12^. The most recent pandemic strain, GII.4 Sydney, accounts for more than half of the HuNoVs identified in acute gastroenteritis cases in children between 2016 and 2020^13^.

The HuNoV genome (∼7.5 kb) encodes three opening reading frames (ORFs)^14^. ORF1 encodes a large precursor polyprotein comprising six nonstructural proteins, including the HuNoV protease. The protease plays a crucial role in viral replication by cleaving the polyprotein into individual nonstructural proteins. Therefore, it has garnered considerable focus as a possible target for developing small-molecule drugs to counter HuNoV infections. HuNoV proteases are cysteine proteases structurally similar to picornavirus 3C-like proteases with a chymotrypsin fold. Their active site features a catalytic triad with histidine-30 (H30) as the catalytic base, cysteine-139 (C139) as the catalytic nucleophile^15^, and glutamic acid-54 (E54) as an acidic residue which orients and stabilizes the conformation of H30^15^. The protease fold comprises two beta barrels: the first contains H30 and E54, while the second beta barrel contains the C139 residue and the substrate binding pocket. The substrate binding pockets S1 to S4 in the HuNoV protease, which accommodates the P1 to P4 residues of the substrate, are in the cavity between the BII-CII loop (beta strands BII and CII, residue 102-117) and the DII, EII, and FII beta strands of the second beta-barrel (**Fig. 1A**).

**Figure 1.**
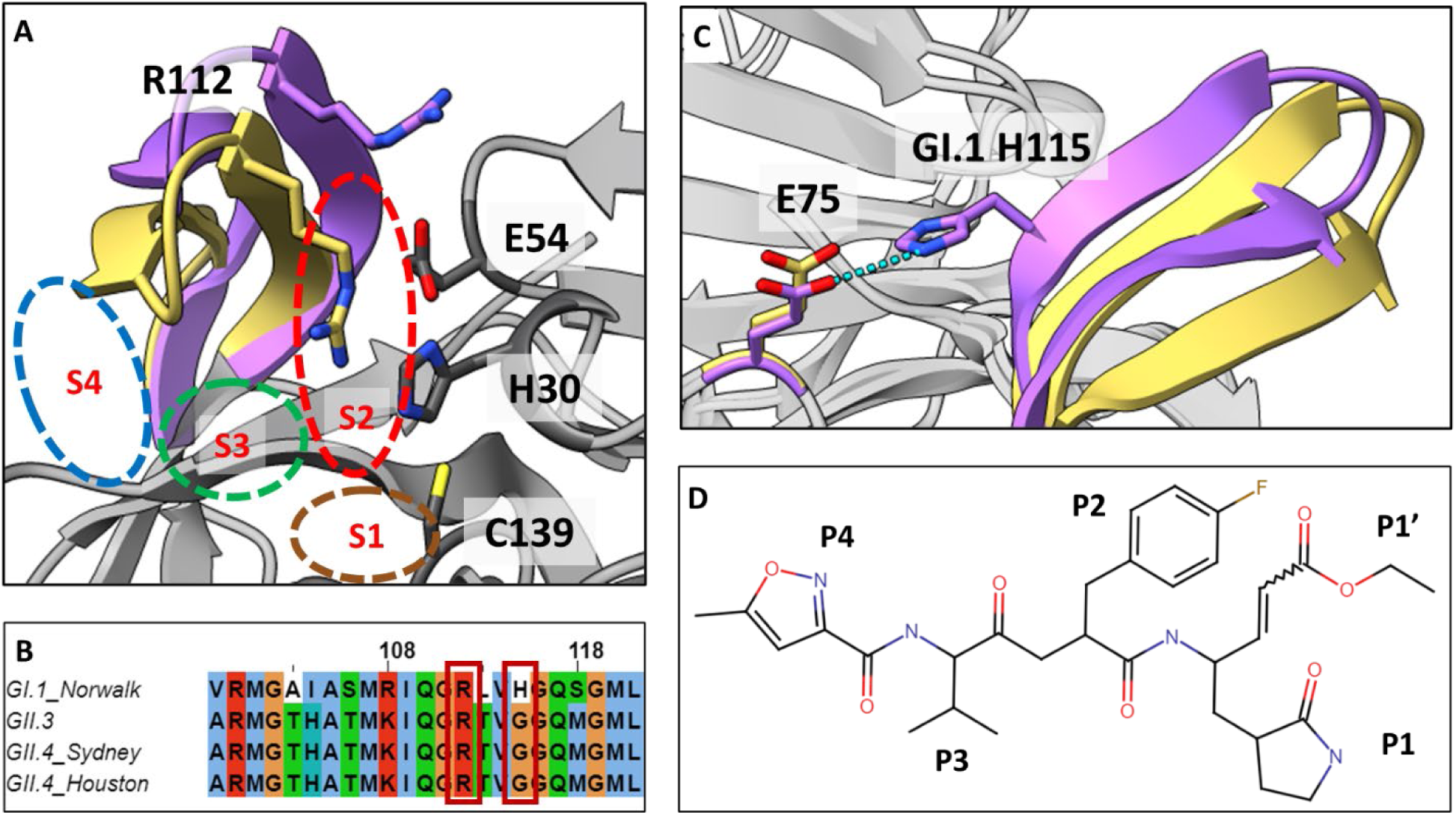
GII proteases exhibit increased flexibility due to sequence changes in the BII-CII loop. **A)** Significant conformational change occurs in BII-CII loop between GI.1 protease (violet) and GII proteases (gold) from residues 102 to 117, while the protein backbone elsewhere remains largely similar. Conserved arginine-112 (R112) points towards the active site in GII protease. **B)** Alignment of BII-CII loop sequence for GI.1 and GII proteases. Conserved arginine-112 and H115G mutation are highlighted. **C)** H115G mutation in GII protease (gold) results in loss of a hydrogen bond that stabilizes the BII-CII loop in the GI.1 protease (violet). **D)** 2D schematic of rupintrivir, the compound chosen to probe the differences between how GI.1 and GII protease respond to inhibitor binding.

GI.1 and GII proteases (GI.1-Pro, GII-Pro) share ∼66% sequence identity. While the sequence changes between these proteases do not alter the overall polypeptide fold, they do cause prominent conformational changes in the substrate binding pockets enclosed by the BII-CII loop, likely necessitated by the changes in the substrate cleavage site of GI.1 and GII polyproteins. In the crystal structure of GI.1-Pro, the BII-CII loop is in the open state (**Fig. 1A**), stabilized by a hydrogen bond **(Fig. 1C**) between histidine-115 (H115) and glutamic acid-75 (E75). In all GII-Pros, H115 is mutated to glycine residue. As observed in the available crystal structures of GII.4-Pro, this H115G mutation results in the loss of this hydrogen bond, causing the BII-CII loop to adopt a closed conformation (**Fig. 1A, gold**). This conformational change in the GII-Pro causes the sidechain of the conserved arginine-112 (R112) to orient closer to the active site^16,17^. The closed conformation of BII-CII loop and the R112 sidechain conformation, as observed in the structures of GII.4-Pro, narrow the S2, S3, and S4 pockets in the apo protease. How these structural changes influence the kinetics and dynamics of substrate and inhibitor interactions generally in GII-Pros and particularly in GII.4-Pros is poorly understood.

The most potent protease inhibitors have been developed primarily against GI.1-Pro, not the proteases of the predominant GII HuNoV strains^18–20^. Compounds tested against both proteases generally show better potency against GI.1-Pro than GII.4-Pro^16,17^. We hypothesize that the lower potency of inhibitors against GII.4-Pro compared to GI.1-Pro is due to the increased flexibility of the BII-CII loop in GII-Pros. Understanding how these structural differences between GI and GII proteases affect inhibitor binding and enzyme kinetics is crucial for developing potent broad-spectrum inhibitors against HuNoV proteases. While the structures of GI and GII-Pros in complex with inhibitors exist, no structures of the same inhibitor bound to GI and GII-Pros are available for a detailed comparative analysis. Existing structures, like the GII.4-Pro in complex with a potent inhibitor^20^, do not yet provide a clear picture of the structural determinants of inhibitor potency in GII.4-Pros. Consequently, how the structural differences between GI and GII-Pros impact substrate binding and inhibitor efficiency remains unclear.

In this study, we determined the crystal structure of GI.1, GII.4, and GII.3-Pros complexed with rupintrivir. We chose rupintrivir, a previously reported nanomolar inhibitor of human rhinovirus 3C protease^21^, after confirming it to be a potent inhibitor of GI.1-Pro^22^, yet simultaneously a poor inhibitor of all GII-Pros tested. The structures reveal that rupintrivir binding elicits only minor conformational changes in GI.1-Pro, whereas a significant BII-CII loop conformational change occurs in GII-Pros. Modeling of rupintrivir into the structures of apo GII proteases reveals widespread steric clashes between rupintrivir and the protease, explaining the need for GII-Pros to undergo an energetically unfavorable BII-CII loop extension to open the S2-S4 pocket and reorienting the R112 to accommodate rupintrivir. We surmise that the contrast in the potencies of rupintrivir against GI.1 and GII-Pros is due to the increased structural flexibility in the BII-CII loop due to H115G mutation. To understand the role of R112 in ligand interaction, we determined the crystal structures of R112A mutant of GII proteases complexed with rupintrivir as well as enzyme and inhibition kinetics. The results indicate that R112 modulates the ligand binding affinity in the active site suggesting a role in the release of cleaved products during polyprotein processing. Our results provide promising starting points for drug development and optimization of lead compounds for both HuNoV GI and GII-Pros.

## RESULTS

### Rupintrivir is highly potent against GI.1-Pro but less potent against various GII-Pros

Rupintrivir has been previously reported to inhibit both GI.1 and GII.4 proteases^22^. Using a competitive FRET protease assay^23^, we show that while rupintrivir inhibits GI.1-Pro very efficiently, it is not very potent against all GII-Pros tested (GII.4 Sydney, GII.4 HOV, GII.3) (**Fig. 2**). The estimated covalent inhibition rate for GI.1-Pro is at least 50-fold higher than for GII-Pros. Given its contrasting inhibition potencies, we chose rupintrivir as a probe to elucidate the structural basis for the differential inhibition potencies against GI and GII-Pros.

**Figure 2.**
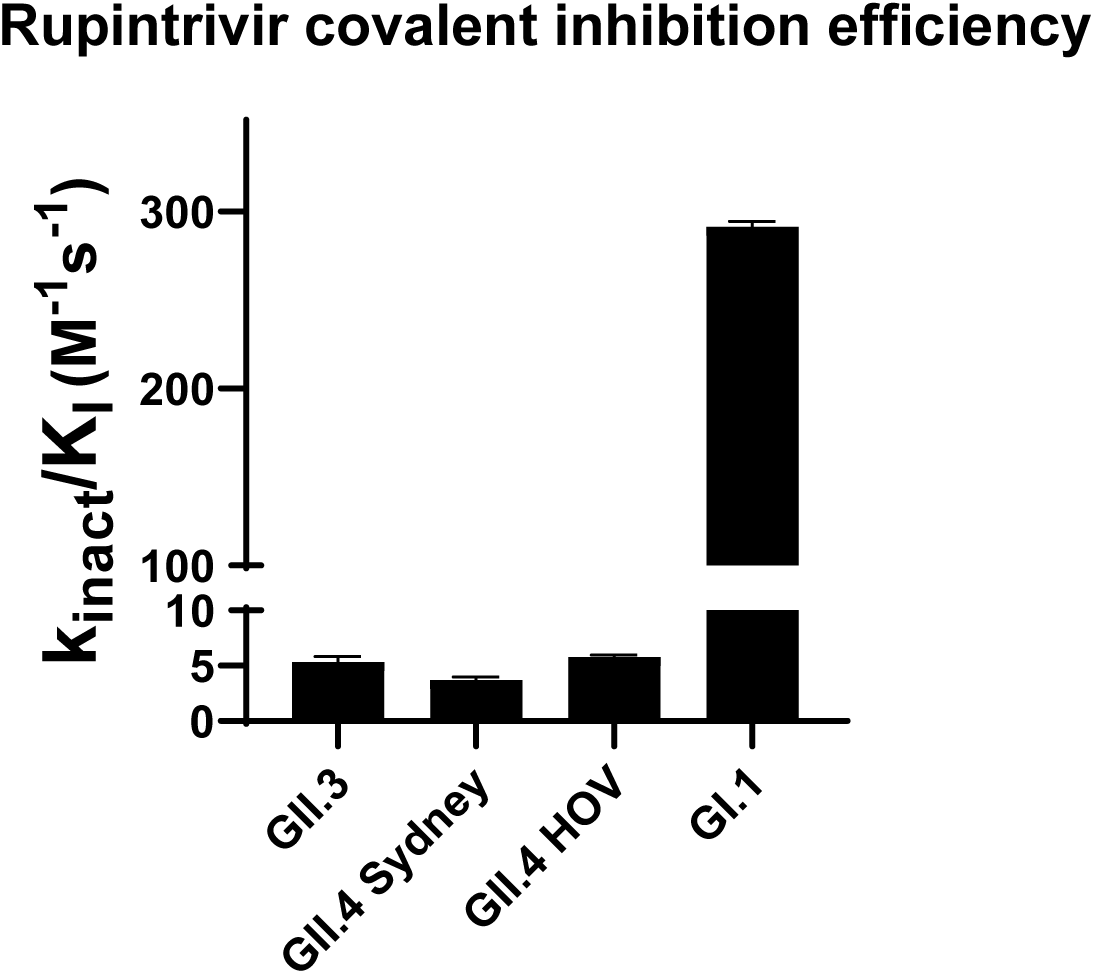
Rupintrivir covalent inhibition efficiencies for GI.1 and GII proteases. Covalent inhibition for HuNoV proteases were measured using continuous, competitive covalent inhibition fluorescent resonance transfer (FRET) assays using fluorogenic peptide substrates corresponding to the p48-p41 cleavage sequence for each genotype.

### Rupintrivir binding requires conformational changes in HuNoV GII-Pros, but not GI.1-Pro

To compare protease-ligand interactions between GI.1 and GII-Pros, we first determined the crystal structures of GI.1 and GII.4 HOV proteases in complex with rupintrivir at 1.7Å and 2.5Å resolution, respectively (**Fig. 3, Table S1**). The GI.1-Pro – rupintrivir complex structure shows excellent pocket complementarity accommodating the P2 to P4 residues of rupintrivir without undergoing any significant conformational changes when compared to previously determined apo GI.1-Pro structure^15^. In contrast, the GII.4-Pro HOV (2002 GII.4 variant) – rupintrivir structure reveals that the BII-CII loop in comparison with published apo structure^16^ undergoes extensive conformational change to accommodate the P3 and P4 residues and the bulky fluorophenylalanine at the P2 position of rupintrivir. However, in this structure, the BII-CII loop extends far past the open conformation observed in apo GI.1-Pro. Considering that part of the loop is involved in crystal packing, the observed conformational change may not entirely be in response to rupintrivir binding as the crystal packing could have also contributed.

**Figure 3.**
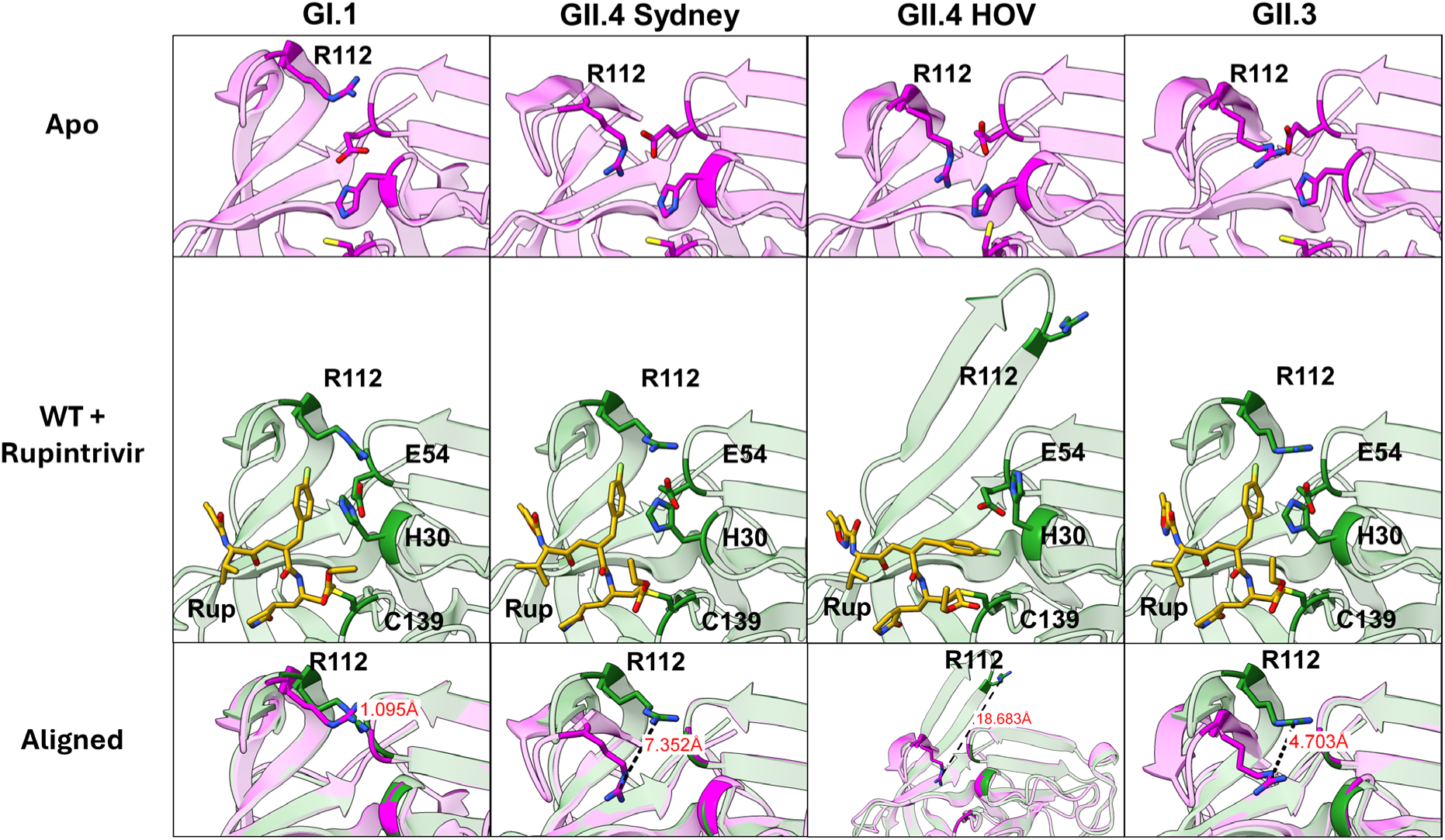
Differential response of GI.1 and GII proteases to rupintrivir binding. GI.1 protease, with BII-CII loop already extended in apo state (top), does not need to undergo conformational change to accommodate rupintrivir. The BII-CII loop is closed in the apo state for GII.4 Sydney, GII.4 HOV and GII.3 proteases (top) and needs to extend to the position equivalent to GI.1 protease to bind rupintrivir (middle). This results in wider conformational change in the arginine-112 sidechain (bottom).

To ascertain that similar conformational changes in the BII-CII loop occur in other GII-Pros in response to rupintrivir binding, we first determined the crystal structures of apo GII.4-Pro Sydney (2012 GII.4 variant) and GII.3-Pro to 2.4Å and 2.6Å, respectively, followed by their crystal structures in complex with rupintrivir to 2.5Å and 2.71Å resolution (**Fig. 3, Table S2**). These structures show that the BII-CII extends from the closed conformation to the same open position observed in GI.1-Pro. The R112 sidechain orients away from the active site, adopting a conformation similar to that observed in GI.1-Pro to accommodate the P2 sidechain of rupintrivir. Unlike the GII.4-Pro HOV-rupintrivir structure, the BII-CII loop is not involved in any crystal contact in these two structures. As a result, the BII-CII loop does not adopt the hyperextended conformation. Also notable is the orientation of the P2 sidechain of rupintrivir. While remaining the same in the GII.4-Pro Sydney and GII.3-Pro structures, this sidechain adopts a different conformation in the GII.4-Pro HOV structure with a hyperextended BII-CII loop.

### BII-CII loop flexibility in GII proteases is necessary for substrate/inhibitor binding

To understand the necessity of the conformational changes in the GII-Pros, we modeled rupintrivir poses into the apo GI.1-Pro and GII-Pro structures. In GI.1-Pro, the modeled rupintrivir molecule introduces minimal steric clashes with the apo protease structure, except for the P1 glutamine mimic, which is readily accommodated through the slight widening of the S1 pocket (**Fig. 4A**). In contrast, the GII-Pro apo structure shows severe clashes with the P2, P3 and P4 groups of the modeled rupintrivir molecule. Isoleucine-109 and methionine-107 clash with rupintrivir’s P3 and P4 groups, respectively, while the P2 of rupintrivir primarily clashes with the R112 sidechain (**Fig. 4B**). Additionally, the fluorine atom of the P2 sidechain causes steric clash with valine-114 (**Fig. 4C**). Thus, loop opening is essential to accommodate the P2, P3 and P4 groups of rupintrivir. However, given the drastic difference between rupintrivir’s potency against GI.1 and GII-Pros, such a loop extension is likely energetically unfavorable.

**Figure 4.**
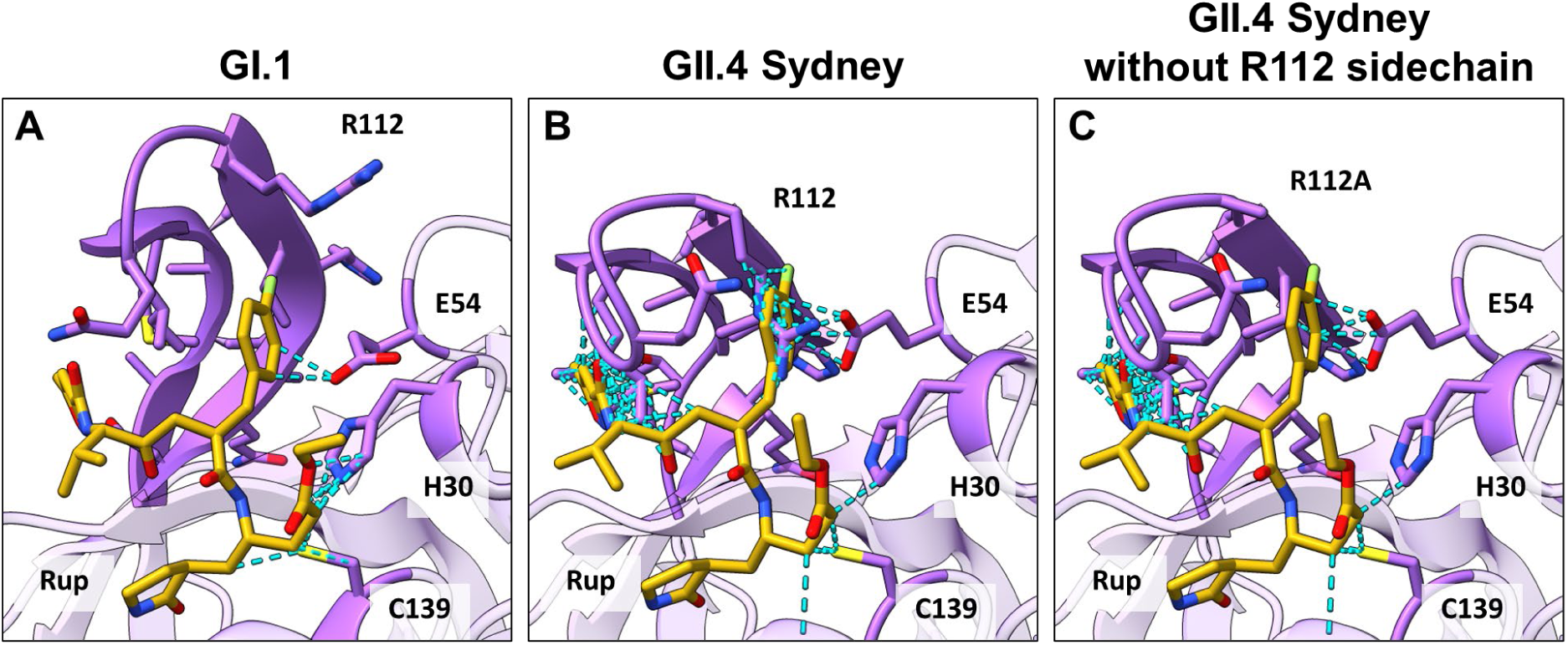
Steric clashes (light blue) between modeled rupintrivir ligand poses and their respective apo protease structures. **A)** GI.1 protease shows minimal steric clash with P2, P3 and P4 residues of rupintrivir. **B)** GII.4 Sydney shows significant steric clashes between BII-CII loop and P3 and P4 of rupintrivir. Additionally, R112 shows steric clash with P2 sidechain of rupintrivir. **C)** Excluding the arginine sidechain, some residual steric clashes remain between valine-104 (also threonine-103, in other GII proteases not shown) and P2 sidechain of rupintrivir. R112 sidechain removed for clarity.

### R112A mutation does not affect rupintrivir binding conformation and its inhibition potency against GII-Pros

Upon rupintrivir binding, not only does the BII-CII loop open in GII-Pros, but the R112 sidechain also extends away from the active site. The P2 sidechain of rupintrivir displaces the R112 sidechain from its relatively stable conformation in the apo protease state. As the R112 apo-protease sidechain conformation and its conformational change upon rupintrivir binding were consistent in all rupintrivir-bound GII-Pro structures, we predicted R112 may interact with the ligand. To investigate the effect of the R112 sidechain on rupintrivir binding and activity, we tested the potency of rupintrivir against the R112A mutants of GI.1, GII.3, GII.4 HOV, and GII.4 Sydney proteases. Considering the covalent inhibition by rupintrivir, we used a previously described procedure to analyze the time-dependent inhibition^24–26^. Surprisingly, rupintrivir shows no significant change in covalent inhibition potency against any of the GII-Pro R112A mutants compared to their respective wild types (WT) while showing lower potency against GI.1-Pro R112A compared to GI.1-Pro WT (**Fig. 5**). This suggests that the presence of the R112 sidechain does not significantly alter rupintrivir’s potency against GII-Pros when measured in our FRET protease assay^16,18^.

**Figure 5.**
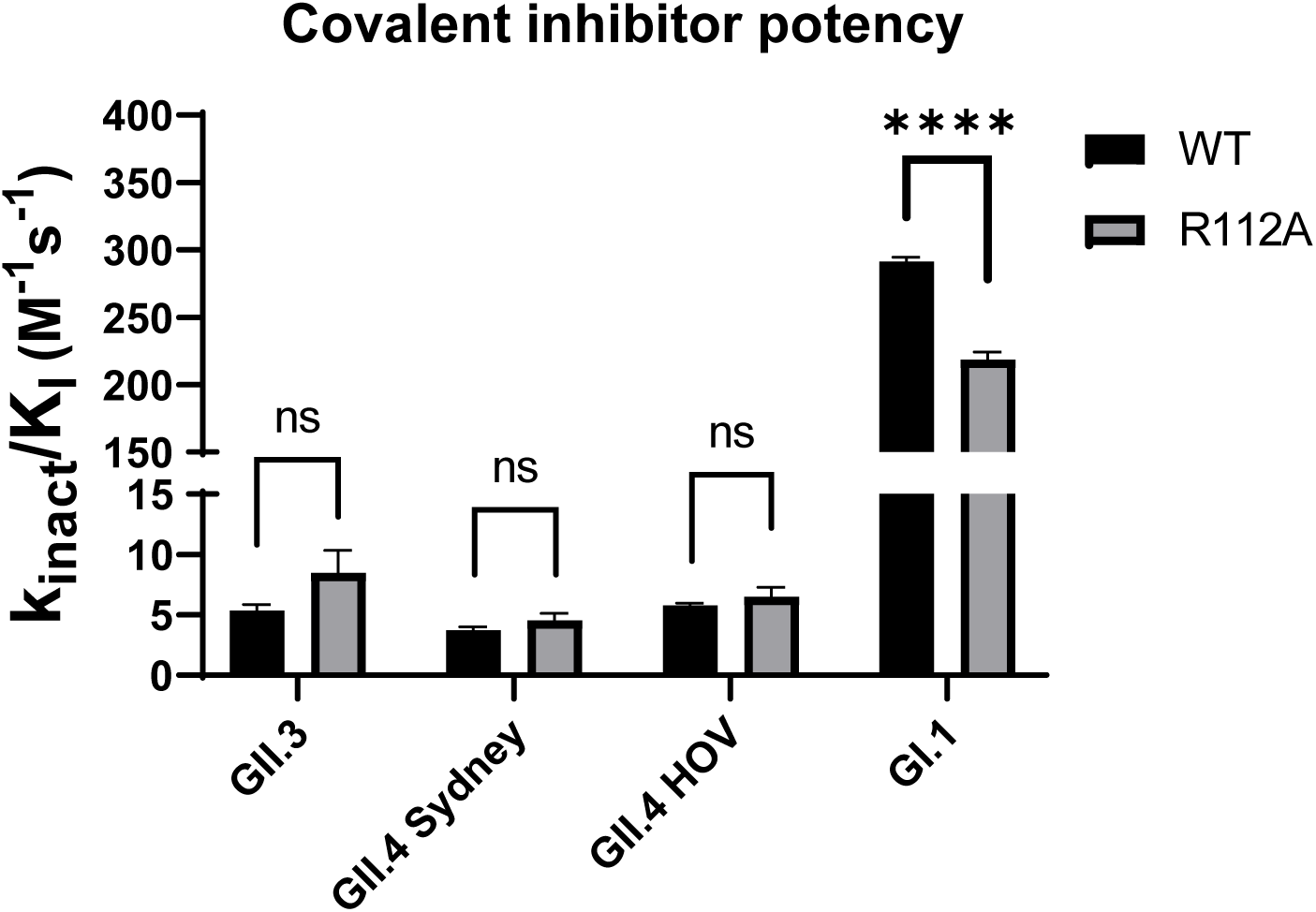
Covalent inhibition efficiency of rupintrivir against WT and R112A of several strains of HuNoV proteases. In all GII strains tested, there were no significant differences in rupintrivir’s covalent inhibition potency against R112A mutant protease compared to their respective WT.

To understand why the R112A mutation does not result in a significant change in potency for rupintrivir despite the obvious steric clashes observed between rupintrivir and the R112 sidechain, we determined the crystal structures of GII.4 Sydney R112A, GII.4 HOV R112A and GII.3 R112A proteases in complex with rupintrivir at 2.4Å, 2.28Å and 2.45Å resolution, respectively. Surprisingly, there is little change in backbone conformation between the respective rupintrivir-bound WT and R112A proteases of all GII strains (**Fig. 6, Table S3**). The BII-CII loops in the R112A protease structures adopt an open conformation even without the R112A sidechain present, with the conformation of the rupintrivir pose remaining the same as in GII.4-Pro Sydney and GII.3-Pro WT structures. In GII.4-Pro HOV R112A, the rupintrivir-bound protease backbone shows the moderately open BII-CII loop position seen in rupintrivir-bound protease structures from other GI and GII strains, indicating that this is the most stable rupintrivir-bound conformation for HuNoV proteases. These highly similar backbones between WT and R112A mutant proteases, especially in the BII-CII loop, further confirm that the main cause for the lower potency of inhibitors against GII-Pro is the steric clash and subsequent unfavorable interaction of the P2, P3, and P4 residues with the BII-CII loop, rather than steric hindrance by the R112 sidechain alone.

**Figure 6.**
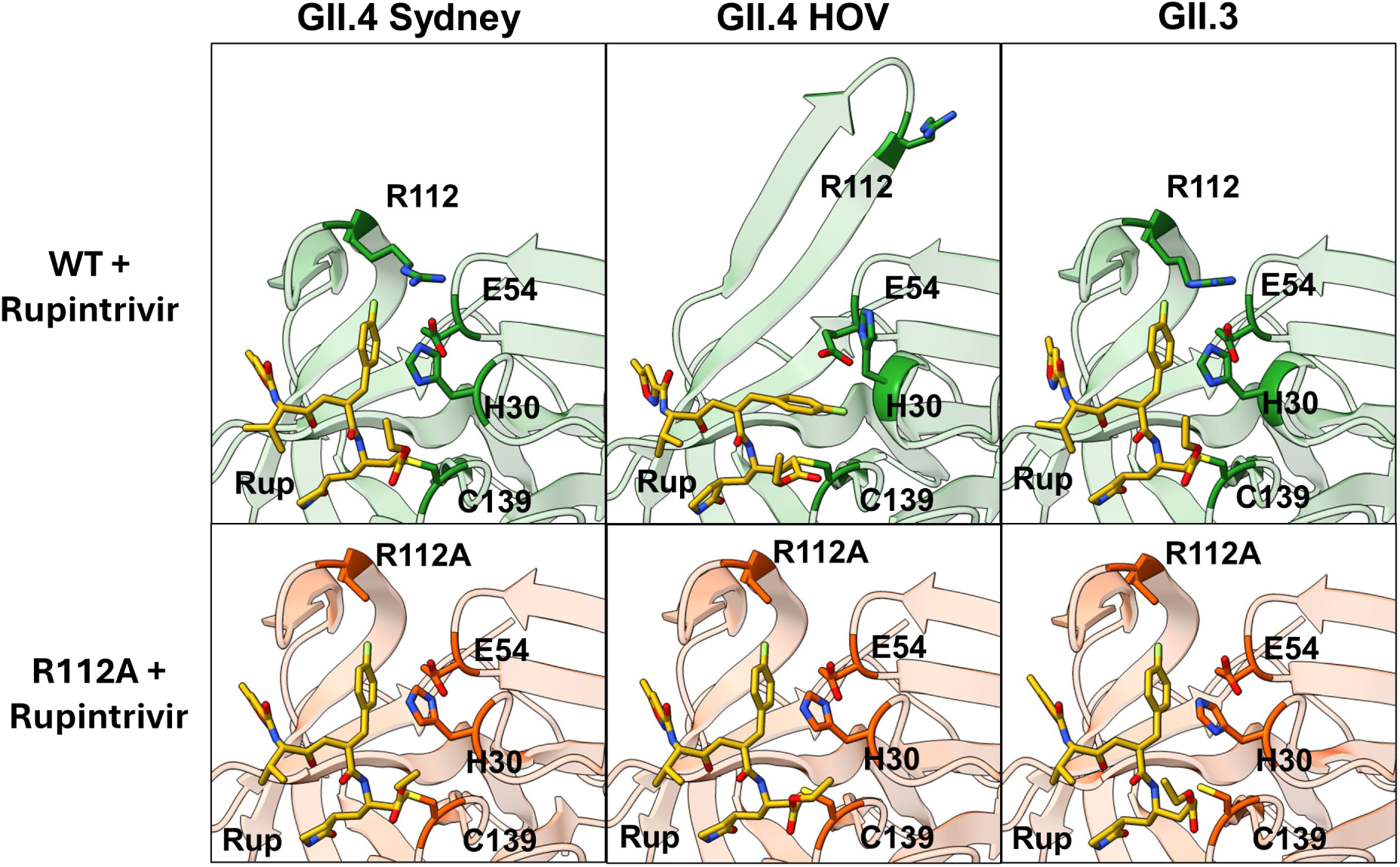
Structural comparison of GII WT and R112A mutant protease – rupintrivir complexes. The protease backbones in R112A mutant protease-rupintrivir structures shows no deviation from protease backbones in their respective WT protease-rupintrivir structures.

### R112 modulates ligand binding affinity with possible implications for product release

To investigate the discrepancy between the conformational shift in the R112 sidechain upon rupintrivir binding and the negligible effect of the R112A mutation on rupintrivir-bound protease structure and rupintrivir potency, we analyzed the enzyme kinetics of the R112A mutants of GII-Pros. The enzyme kinetics results (**Fig. 7**) for WT and R112A proteases of GI and GII strains revealed a generally decreasing enzyme turnover rate (k_cat_) upon R112A mutation. Additionally, we observed increasing substrate affinity (decreasing Michaelis constant, K_M_) for all proteases, except for GII.3, which already has the lowest Michaelis constant. Together, this results in a significant decrease in protease catalytic efficiency (k_cat_/K_M_) for GII proteases but no substantial change in GI.1 protease. These data suggest that R112 plays a vital role in enzyme turnover and possibly in direct displacement of the released cleavage product and any ligand bound in the active site. We hypothesize that the R112 sidechain interacts broadly with any ligand in the active site by sterically restricting the P2 sidechain.

**Figure 7.**
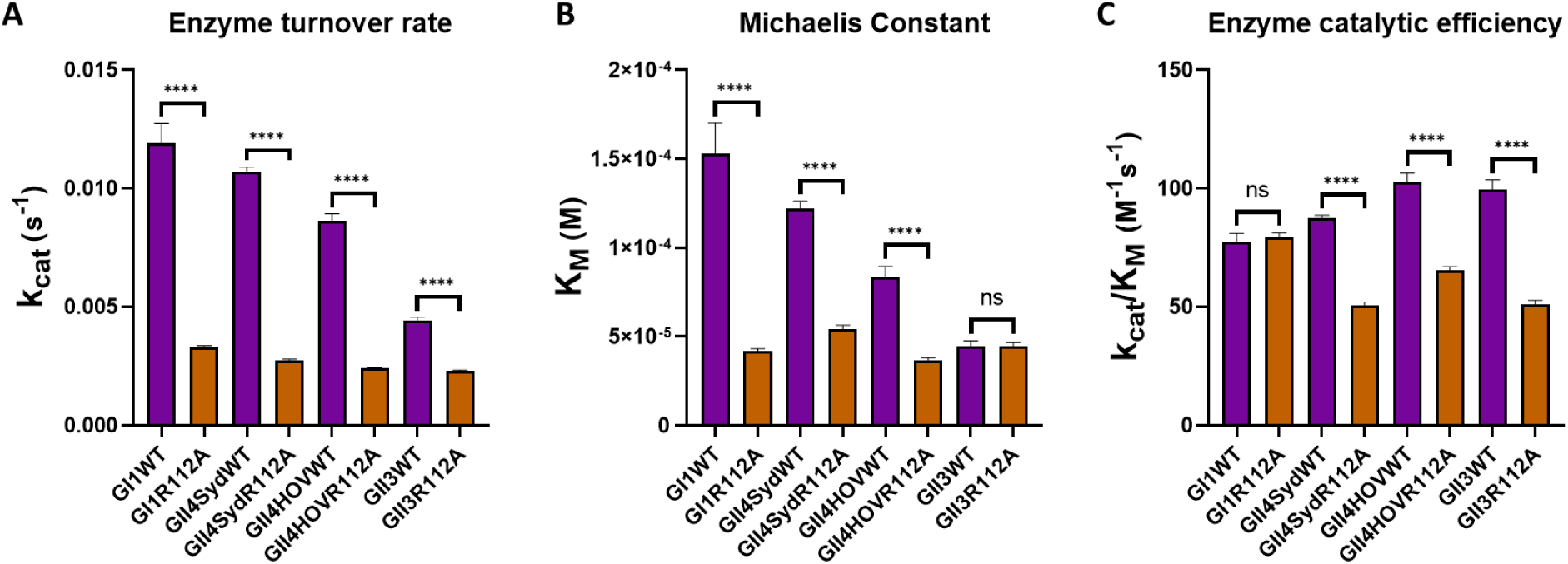
Enzyme kinetics characteristics of WT and R112A mutant GI.1 and GII proteases. **A)** R112A mutation decreases enzyme turnover rate for HuNoV proteases. **B)** For all proteases tested except GII.3 protease, Michaelis constant decreases with the R112A mutation, indicating improved substrate affinity. **C)** For GI.1 protease, R112A mutation does not significantly change catalytic efficiency. For GII proteases, R112A mutation decreases the protease catalytic efficiency.

## DISCUSSION

The genetic divergence from GI to GII HuNoVs resulted in significant changes in the HuNoV protease sequence and structure, as well as substrate sequence and recognition. Consequently, inhibitors developed against GI.1-Pro are generally much less effective against GII-Pros. Previous studies have identified closed BII-CII loop and placement of R112 in the active site as significant structural changes between GI.1-Pro and GII-Pros^16,27^. However, the mechanism by which these changes adversely affect inhibitor potency in GII-Pros is poorly understood. Our studies here show that the substrate binding pocket opens through BII-CII loop extension to bind inhibitors like rupintrivir, an ostensibly energetically unfavorable process enabled only by the conformational flexibility in the BII-CII loop of GII-Pros.

### Structural basis for reduced inhibitor potency against GII-Pros

Our FRET inhibition data analysis reveals that rupintrivir is almost two orders of magnitude less potent against GII-Pros than GI.1-Pro. In GII-Pros, but not GI.1-Pro, the BII-CII loop switches from a closed state to an open state to accommodate rupintrivir. The R112 sidechain also changes from a stable conformation in the active site to a less stable conformation away from the active site. However, our assays with R112A mutants of GII-Pro reveal that the R112A mutation does not significantly increase the potency of rupintrivir against GII-Pros. Our structures of GII R112A proteases in complex with rupintrivir further show no significant changes in the protease backbone between rupintrivir-bound WT and R112A proteases. These results suggest that the differential placement of R112 in GII-Pros is not solely responsible for the observed potency difference.

Therefore, the interaction between rupintrivir and the entire BII-CII loop dictates the reduced potency of rupintrivir. Our modeling shows that the steric clash between the apo GII-Pro and P2-P4 of rupintrivir forces the BII-CII loop to open and adopt an energetically unfavorable raised conformation. Therefore, the binding energy between the BII-CII loop and rupintrivir (any substrate in general) in GII-Pros must counterbalance such an unfavorable conformation. In GI.1-Pro, the H115-E75 hydrogen bond is positioned in the middle of the BII-CII loop, effectively stabilizing it. In GII-Pros, the BII-CII loop is stabilized only by a hydrogen bond between histidine-104 and alanine-79, located at the bottom of the loop. While rupintrivir makes backbone hydrogen bond contacts with the inflexible portion of the binding pocket, i.e., alanine-158 and alanine-160, the molecule relies mainly on van der Waals interactions to bind the BII-CII loop. We hypothesize that the BII-CII loop opening cannot be effectively stabilized by these interactions alone, resulting in the observed drop in potency for rupintrivir from GI.1-Pro to GII-Pros.

### Differential conformational dynamics of inhibitor binding in GI.1 and GII-Pros

Interestingly, in the recently published structure of GII.4-Pro Sydney in complex with NV-004, a potent aldehyde-based inhibitor of SARS-CoV-2 main protease with HuNoV protease inhibition activity^28,29^ (**Fig. S1**), the BII-CII loop exists in a different conformation compared to our rupintrivir bound GII-Pro structures. In the NV-004 bound structure, the loop conformation results in a narrower S2 pocket and a wider S4 pocket, suggesting that in exchange for stability, the BII-CII loop matches its conformation to the bound ligand to attain the most favorable conformation. These comparative analyses affirm that the S2-S4 substrate binding pockets are more flexible in GII-Pros than in GI.1-Pro, in which the S2 pocket is constrained to an open position by the H115-E75 hydrogen bond. It is unclear what advantage the transition to flexible substrate recognition confers to the GII-Pros. The flexible BII-CII loop in GII-Pros perhaps offers an evolutionary advantage for accommodating sidechain variations in the substrate cleavage sites and based on our kinetic analysis (**Fig. 7**), the flexibility also could result in increased affinity for the substrates and higher catalytic efficiency.

### R112 sidechain likely plays a role in displacing substrates from the active site

From our rupintrivir bound GII-Pro WT structures, we conjectured that removing the steric clash due to R112 by R112A mutation would improve the inhibitor potency in GII-Pros. However, we observed that the R112A mutants of GII-Pro are enzymatically less efficient than their respective WT, and surprisingly, the R112A mutation in these proteases has minimal effect on rupintrivir inhibition. Our enzymatic assays showed a significant reduction in enzyme turnover rate (k_cat_) with R112A mutants of all proteases tested, including GI.1-Pros. In addition, except in the case of GII.3-Pro, there was a reduction in the Michaelis constant (K_M_). In GII-Pros, but not in GI.1-Pro, the change in enzyme turnover rate outweighs the change in Michaelis constant, resulting in a significant drop in protease catalytic efficiency. These results suggest that the R112A mutation increases substrate, ligand, and inhibitor affinities. Although the reduced K_M_ value suggests an enhanced affinity for the substrate, we cannot rule out that the lower K_M_ is instead due to a slower and rate-limiting deacylation of the covalent thioester intermediate in the protease reaction mechanism, which is also consistent with reduced k_cat_ and K_M_ values^30,31^.

While the rate-limiting step in the catalytic cycle of HuNoV protease is not known, insufficient steric control of bound substrate has been implicated in attenuating enzymatic activity by impairing product release and shifting the rate-limiting step to product release^32^. In UDP-galactopyranose mutase, this occurs during catalysis of a smaller substrate compared to the natural substrate or when the bulkier side chain, such as tryptophan, interacting with the substrate, is mutated to a smaller side chain such as alanine. Similarly, we posit that the R112A mutation likely hinders ligand displacement by the protease, resulting in increased affinity to the protease for all ligands. We further hypothesize that the R112 sidechain primarily interacts with the P2 sidechain. Considering the similar bulkiness of the P2 sidechain of our FRET substrate and rupintrivir, the R112A mutation does not significantly shift the binding competition between rupintrivir and the substrate. As a result, we observe no changes in rupintrivir potency against the protease. These results imply that with its flexible sidechain, R112 may displace bound ligands, including cleaved products and inhibitors, by acting on P2-equivalent moieties positioned in the S2 pocket.

Flexible sidechains near the substrate binding pocket have been proposed to play a role in product release in other proteases. Asparagine-142 of SARS-CoV-2 main protease is dynamic and flexible, facilitating its role in post-cleavage product release^33^. Similarly, several residues in Dengue Virus protease are thought to be involved in product release, as they affect activity in steady-state conditions but not in single-turnover conditions^34^. Of note, in 3C-like proteases structurally related to HuNoV protease, we observe very similar charged, flexible residues in R112 position, arginine-130 in poliovirus protease, lysine-130 in enterovirus A71 protease, and asparagine-130 in human rhinovirus protease^21,35,36^.

### Implication for the development of broad-spectrum inhibitors against HuNoV proteases

Our studies strongly suggest that R112 sidechain likely influences product release through steric hindrance and is an important determinant of substrate and inhibitor binding. Since R112 primarily interacts with the P2 position, optimizing the P2 sidechain in peptidomimetic inhibitors is a potential avenue to improve potency broadly across HuNoV proteases. Smaller sidechains than leucine and phenylalanine, as found in the P2 position of the HuNoV substrates, may be more resistant to displacement by the R112 sidechain and might result in increased binding affinity of the inhibitor to the protease. It is still unclear how the changes in the S4 pocket, as noted above affect substrate and inhibitor binding in GII-Pros. Based on GII.4-Pro Sydney structures with different inhibitors, we hypothesize that the P2 and P4 sidechains being simultaneously bulky would compromise inhibitor affinity. The P3 and P4 positions, therefore, likely have some influence on loop opening. Diverse and novel chemotypes replacing the P3 and P4 positions have been demonstrated for HIV protease and GI.1-Pro^29,37^, with one such inhibitor being effective on GII.4-Pro^20^. These studies indicate the S4 pocket can accommodate more diverse binding moieties than previously assumed. The P3 sidechain is not directly involved in binding the protease and instead torsionally constrains the P4 and P2 sidechains into the correct positions. Replacing the P3 with nonpeptidyl groups is also viable^37^. Finding novel and effective chemotypes to replace the peptidyl P3-P4 groups partially or entirely may be a necessary step towards finding potent and broad spectrum HuNoV protease inhibitors.

In summary, our studies identified BII-CII loop interactions as important determinants for inhibitor potency in GII proteases. Though the BII-CII loop opening is necessary for binding inhibitors, the loop opening is unfavorable and needs to be stabilized with inhibitor interactions. The loop can flexibly adopt different conformations when binding different inhibitors; however, bulky inhibitor sidechains likely result in unfavorable binding and lower inhibitor potency. Furthermore, we found that the R112 residue negatively affects substrate affinity in GII proteases and theorize that its role is to release cleaved products from the active site. Our inhibition assays with R112A GII protease mutants reveal that R112 also adversely affects rupintrivir’s affinity. This indicates that the highly conserved R112 may play a critical role in displacing bound ligands and is necessary for efficient polyprotein processing. The structural basis of how GI and GII HuNoV proteases differentially respond to inhibitor binding and affect the inhibition potencies we have described here can be useful for the design of more potent broad-spectrum HuNoV proteases.

## MATERIALS AND METHODS

### Expression and purification of HuNoV proteases

GI.1-Pro and GII.4-Pro Houston were cloned into pET-46b(+) vectors with a thrombin site added after the enterokinase site. GII.4-Pro Sydney (AFV08794.1) and GII.3-Pro (BAG30938.1) were cloned into a pET-based vector with a 6xHis-TELSAM fusion tag^38^ followed by a 3C protease cleavage site. Transformed XJb(DE3) *E. coli* (Zymo Research) grown overnight at 30°C in Terrific Broth (TB) supplemented with the appropriate antibiotics and 1.5% glucose were used to inoculate TB supplemented with antibiotics, 0.4% glycerol, 0.05% glucose and 2mM MgCl_2_ at 37°C. The cultures were grown to an optical density of 0.7 before induction with 0.4mM IPTG. The temperature is reduced to 18°C and the culture is grown overnight for 16 hours before collection and centrifugation at 3,500g for 30 minutes. The pellet is resuspended in lysis buffer (20mM HEPES, 500mM NaCl, 20mM imidazole, 1mM TCEP, pH 7.5) supplemented with GENIUS^TM^ nuclease (ACROBiosystems). The suspension is lysed with the LM-20 Microfluidizer (Microfluidics) at 18,000 PSI, and the lysate is centrifuged at 35,000g for 30 minutes. The supernatant is incubated with Ni-NTA resin pre-equilibrated in lysis buffer at 4°C with rocking for 1 hour. The resin is then centrifuged and decanted, then washed 4 times with wash buffer (20mM HEPES, 1M NaCl, 30mM imidazole, 1mM TCEP, pH 7.5) before transferring to a glass Econo-Column (Bio-Rad). The resin is washed once with lysis buffer before elution with elution buffer (20mM HEPES, 150mM NaCl, 300mM imidazole, 1mM TCEP, pH 7.5) in elution with 6 column volumes. When used, 3C protease (pET-NT*-HRV3CP was a gift from Gottfried Otting (Addgene plasmid #162795; http://n2t.net/addgene:162795; RRID:Addgene_162795) was added to the elution at 1:50 3C protease to His-tagged protein mass:mass ratio, and human alpha-thrombin (Haematologic Tech) was added to the elution at 1:2000 v/v ratio. The mixture was transferred to SnakeSkin dialysis tubing (7K MWCO, ThermoFisher) and dialyzed against dialysis buffer (10mM HEPES, 100mM NaCl, 15mM imidazole, 1mM TCEP, pH 8.0) overnight with gentle stirring. The mixture was incubated with Ni-NTA resin pre-equilibrated with size exclusion chromatography buffer (10mM HEPES, 50mM NaCl, 1mM TCEP, pH 8.0) for 80 minutes at 4°C. The flowthrough was collected and concentrated to 5ml before injection on a HiLoad 16/60 Superdex 75 pg column (Cytiva) equilibrated in size exclusion chromatography buffer. The fractions containing the protease were pooled. The sample for crystallography is concentrated and used or flash-frozen in liquid nitrogen immediately, while samples to be used in enzymatic assays were supplemented with 50% glycerol to 10% final glycerol concentration, then concentrated to around 10 mg/ml, flash-frozen in liquid nitrogen and stored at −80°C.

### Enzymatic and time-dependent inhibition assay

The activity of the proteases was measured using a fluorescent resonance transfer assay as described previously. Fluorogenic substrate peptides (GI.1 substrate: Glu(EDANS)- PDFHLQGPEDLA-Lys(Dabcyl), GII substrate: Glu(EDANS)-GDYELQGPEDLA-Lys(Dabcyl), corresponding to the cleavage site between p48 and p41 subunits in the polyprotein) were synthesized by GenScript USA Inc. For enzyme kinetics assays, 1.2mM substrate solutions in 100% DMSO was diluted with assay buffer (10mM HEPES, 30% glycerol, 10mM DTT, 0.1% CHAPS, pH 8.0) to 256µM and further serially diluted two-fold with assay buffer down to 1µM. A 2µM protease solution was prepared by diluting a stock protease solution in assay buffer. 50µl of substrate solution was then added to 50µl of a 2µM protease solution pre-dispensed in 96-well black NBS plates (Corning 3991), briefly mixed with a multichannel pipette before shaking for 90 seconds at 37°C, 1000RPM in the FlexStation 3 multimode plate reader (Molecular Devices). Assays were run for 2 hours at 37°C with 90-second measurement intervals (excitation 340nm, emission 490nm, filter 475nm,). The product concentration was calculated from the relative fluorescence units (RFU) using a standard curve. Initial velocities were calculated in GraphPad Prism 8 software (GraphPad Software Inc.). Nonlinear regression analysis in GraphPad Prism 8 was used to fit Michaelis constants (*K*_M_), the catalytic constant (*k*_cat_), and calculate the catalytic efficiency (*k*_cat_/*K*_M_).

For inhibition assays, 1.2mM substrate solutions in DMSO was diluted with assay buffer to 100µM. Rupintrivir stock solution (40mM in DMSO) was diluted with DMSO to the final concentration ranges of 0.225-3.6mM (GII-Pro WT), 0.112-1.8mM (GII-Pro R112A), and 6.25µM-50µM (GI.1-Pro, WT and R112A), respectively. The substrate solution was mixed with rupintrivir solution at 5:1 v/v ratio, and 60ul of this mixture was dispensed into each well of the 96 well plate (Corning 3991). 40ul of a 1µM protein solution was dispensed into each well and mixed briefly before the plate is sealed with ClearSeal film (Hampton Research). The plate is shaken for 90 seconds at 37°C in the FlexStation 3, before assays were run for 4 hours at 37°C with 90-second measurement intervals. Relative fluorescence unit (F) versus time (T) curves were fitted using the *scipy.optimize*^39^ Python library using Equation 1, where V_s_ is the steady-state velocity, V_0_ is the initial velocity, k_obs_ is the observed covalent inhibition rate, and F_0_ is the offset. The initial parameters for F_0_, V_0_, V_s_ and k_obs_ were estimated using the *differential_evolution* function in *scipy.optimize*. Then, the *curve_fit* function in scipy optimize was used to fit F_0_, V_0_, V_s_ and k_obs_ using the bounded ‘trf’ option.

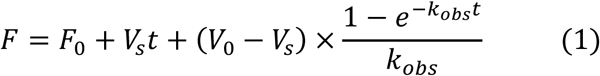

The obtained k_obs_ values were imported into GraphPad Prism 8; k_inact_ and K_I_^app^ for GII-Pros were fit using nonlinear regression using equation 2, where [I] is the inhibitor concentration and k_ctrl_ is the observed degradation rate in the control samples with only protease and substrate.

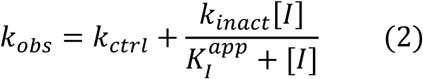

For GI.1-Pro, as the reaction occurs in inhibitor tight binding conditions, Equation 3 was used instead, and k_inact_/K_I_^app^ = k_chem_^app^ approximation was used.

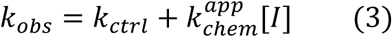

The correction for substrate competition was done with Equation 4, where [S] is the substrate concentration and K_M_ is the Michaelis constant for the respective protease and substrate pair.

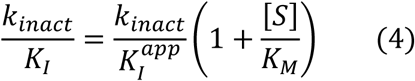

### Crystallization, data collection, and refinement

GII.4-Pro Sydney and GII.3-Pro were concentrated to 18 mg/ml and 12 mg/ml, respectively, before setting up for crystallization trials. 50µl of crystallization conditions were dispensed into 96-well Intelli-plate crystallization plates (Art Robbins), and the Mosquito liquid handler (SPT Labtech) was used to add 0.2µl protein and 0.2µl of crystallization solution to sitting drop wells of the same plate. The plate was then sealed with Crystal Clear Sealing Film (Hampton Research).

To obtain rupintrivir-bound crystals, 200µl of 40mM rupintrivir (MilliporeSigma) dissolved in 100% DMSO was diluted in between 10 and 15ml of SEC buffer (10mM HEPES, 50mM NaCl, 1mM TCEP, pH 8.0) and mixed gently and thoroughly. Then, protease solution at 10mg/ml concentration or higher were added to a final protein:rupintrivir ratio of 1:15 or 1:20 and left to mix by gentle rocking at 4°C overnight. For GII.4-Pro Sydney R112A, the SEC buffer was supplemented with 1M potassium iodide to a final concentration of 200mM. After overnight incubation, the protein was concentrated with an Amicon Ultra-4 filter to a final absorbance at 280nm between 6 and 8 and used for crystallization in sitting drop plates with 50µl reservoir solution and 0.2µl protein plus 0.2µl reservoir solution drops.

All crystals formed between 3 and 7 days after setting up. Crystals were harvested and immediately flash-frozen in liquid nitrogen without additional cryoprotection, except for GII.4-Pro HOV WT – rupintrivir and GII.3-Pro R112A – rupintrivir, for which 20% glycerol was used for cryoprotection. Crystallization conditions for all final crystals used for X-ray diffraction data collection using the synchrotron beamline facilities at ALS, SSRL, or APS are summarized in **Table S4**.

X-ray crystal diffraction images of all the crystals were processed with the *xia2* package using either the dials or XDS pipelines^40–45^. Molecular replacement (PHASER^46^), refinement (phenix.refine^47–51^) and ligand restraints (ReadySet) was done in PHENIX^51^, with model building done in COOT^52^. GII.4 Houston protease apo structure (PDB ID: 6NIR) was used as template for molecular replacement for GII.4-Pro Houston, GII.4-Pro Sydney and GII.3-Pro structures; GI.1-Pro apo structure (PDB ID: 2FYQ) was used as template for molecular replacement for GI.1-Pro – rupintrivir complex structure. Data collection and refinement statistics and crystallization conditions can be found in supplementary materials. Figures were prepared using ChimeraX^53–55^. Sequence alignments were carried out using Jalview^56^.

## ACKNOWLEDGEMENTS

This work was supported by the National Institutes of Health (NIH) grant P01 AI057788 (to MKE, Robert Atmar, and BVVP) and a grant from the Robert Welch Foundation (Q-1279 to BVVP). Beamlines 5.0.1, 5.0.2, 8.2.1 and 8.2.2 of the Advanced Light Source, a DOE Office of Science User Facility under Contract No. DE-AC02-05CH11231, are supported in part by the ALS-ENABLE program funded by the NIH, National Institute of General Medical Sciences, grant P30 GM124169-01. Use of the Stanford Synchrotron Radiation Lightsource, SLAC National Accelerator Laboratory, is supported by the US Department of Energy, Office of Science, Office of Basic Energy Sciences under Contract No. DE-AC02-76SF00515. The SSRL Structural Molecular Biology Program is supported by the DOE Office of Biological and Environmental Research and by the National Institutes of Health, National Institute of General Medical Sciences (P30GM133894). The contents of this publication are solely the responsibility of the authors and do not necessarily represent the official views of NIGMS or NIH. This research used resources of the Advanced Photon Source, a US Department of Energy (DOE) Office of Science user facility operated for the DOE Office of Science by Argonne National Laboratory under Contract No. DE-AC02-06CH11357.

