## Supplementary material for "Conformational flexibility is a critical factor in designing broad-spectrum human norovirus protease inhibitors": Supplemantary Figure S1 and Tables S1-S4

**This file includes:**

Supplementary Figure S1

Tables S1 to S4

**Supplementary Figure S1**

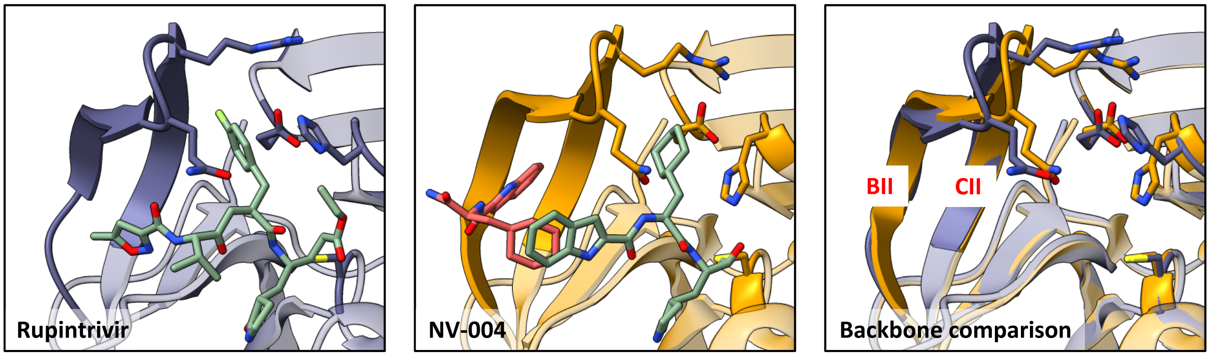


**Supplementary Figure S1. Comparison of GII.4 Sydney protease-rupintrivir and GII.4 Sydney-NV-004 structures. Left:** GII.4 Sydney is covalently linked to a single molecule of rupintrivir. **Middle:** GII.4 Sydney is linked to one molecule of NV-004 (light blue) while binding another NV-004 molecule (pink) in the S4 pocket. **Right:** The CII strand of Sydney-NV-004 structure (gold) is less extended than that of Sydney-rupintrivir structure (indigo), while BII-CII of Sydney-NV-004 is more extended than that of Sydney-rupintrivir structure, likely to match the ligands present in their respective S2 and S4 pockets.

**Table S1: Data collection and refinement statistics for GI.1-Pro-rupintrivir and GII.4-Pro HOV-rupintrivr structures**

| | GI1_WT_Rupintrivir | HOV_WT_Rupintrivir |
|---|---|---|
| PDB ID | 9D9Y | 9DA7 |
| Wavelength | | |
| Resolution range | 52.98 - 1.7 (1.75 - 1.7) | 48.99 - 2.5 (3.15 - 2.5) |
| Space group | P 43 21 2 | P 31 2 1 |
| Unit cell | 118.464 118.464 66.571 90 90 90 | 56.569 56.569 95.064 90 90 120 |
| Total reflections | 457948 (37429) | 79537 (40190) |
| Unique reflections | 52590 (4175) | 6478 (3176) |
| Multiplicity | 8.7 (9.0) | 12.3 (12.7) |
| Completeness (%) | 99.95 (99.91) | 99.52 (99.78) |
| Mean I/sigma(I) | 10.13 (1.58) | 16.98 (11.91) |
| Wilson B-factor | 23.7 | 43 |
| R-merge | 0.1008 (1.673) | 0.08316 (0.134) |
| R-meas | 0.1072 (1.772) | 0.08694 (0.1397) |
| R-pim | 0.03572 (0.5778) | 0.02476 (0.03898) |
| CC1/2 | 0.998 (0.779) | 0.997 (0.996) |
| CC* | 1 (0.936) | 0.999 (0.999) |
| Reflections used in refinement | 52505 (4294) | 6438 (3169) |
| Reflections used for R-free | 1701 (138) | 326 (160) |
| R-work | 0.1855 (0.3060) | 0.2515 (0.3204) |
| R-free | 0.2075 (0.2913) | 0.2914 (0.3896) |
| CC(work) | | |
| CC(free) | | |
| Number of non-hydrogen atoms | 2776 | 1261 |
| macromolecules | 2458 | 1201 |
| ligands | 86 | 43 |
| solvent | 232 | 17 |
| Protein residues | 328 | 159 |
| Nucleic acid bases | | |
| RMS(bonds) | 0.008 | 0.005 |
| RMS(angles) | 1 | 1.18 |
| Ramachandran favored (%) | 98.44 | 97.42 |
| Ramachandran allowed (%) | 1.56 | 2.58 |
| Ramachandran outliers (%) | 0 | 0 |
| Rotamer outliers (%) | 0.38 | 0.78 |
| Clashscore | 2.55 | 0.8 |
| Average B-factor | 28.28 | 46.21 |

| | | |
|---|---|---|
| macromolecules | 27.3 | 45.97 |
| ligands | 26.41 | 54.67 |
| solvent | 39.45 | 41.41 |
| Number of TLS groups | | |

**Table S2: Data collection and refinement statistics for GII.3-Pro and GII.4-Pro Sydney apo and rupintrivir complex structures**

| | GII3_WT_Apo | GII4_Sydney_Apo | GII3_WT_Rupintrivir | Sydney_WT_Rupintrivir |
|---|---|---|---|---|
| PDB ID | 9DF5 | 9DEY | 9DAP | 9DAJ |
| Wavelength | | | | |
| Resolution range | 63.04 - 2.6 (2.7 - 2.6) | 60.54 - 2.4 (2.5 - 2.4) | 48.66 - 2.71 (3.41 - 2.71) | 48.86 - 2.5 (2.86 - 2.5) |
| Space group | P 43 | C 2 2 21 | P 32 2 1 | P 32 2 1 |
| Unit cell | 78.526 78.526 105.739 90 90 90 | 112.21 158.9 93.5 90 90 90 | 97.329 97.329 43.77 90 90 120 | 97.722 97.722 44.62 90 90 120 |
| Total reflections | 190806 (21594) | 142240 (15873) | 334419 (166547) | 81116 (26627) |
| Unique reflections | 19982 (2177) | 31740 (3522) | 6699 (3293) | 8839 (2882) |
| Multiplicity | 9.5 (9.9) | 4.5 (4.5) | 49.9 (50.6) | 9.2 (9.2) |
| Completeness (%) | 99.95 (100.00) | 92.94 (65.72) | 99.78 (99.67) | 99.53 (98.98) |
| Mean I/sigma(I) | 15.77 (2.39) | 4.53 (2.02) | 5.28 (3.42) | 15.61 (4.76) |
| Wilson B-factor | 57.79 | 22.86 | 50.06 | 42.87 |
| R-merge | 0.09754 (0.9054) | 0.1795 (0.4681) | 0.8046 (1.102) | 0.08504 (0.3243) |
| R-meas | 0.1032 (0.9549) | 0.2084 (0.5453) | 0.8121 (1.112) | 0.09046 (0.3454) |
| R-pim | 0.03323 (0.3016) | 0.1019 (0.2695) | 0.1087 (0.1443) | 0.03033 (0.1168) |
| CC1/2 | 0.998 (0.784) | 0.797 (0.477) | 0.953 (0.906) | 0.997 (0.961) |
| CC* | 1 (0.937) | 0.942 (0.804) | 0.988 (0.975) | 0.999 (0.99) |
| Reflections used in refinement | 19778 (2216) | 30711 (2391) | 6684 (3282) | 8664 (2815) |
| Reflections used for R-free | 1217 (136) | 1207 (93) | 318 (165) | 440 (137) |
| R-work | 0.2724 | 0.2253 (0.3059) | 0.1862 (0.2594) | 0.2290 (0.2968) |
| R-free | 0.2876 | 0.2594 (0.3326) | 0.2402 (0.2973) | 0.2786 (0.3192) |
| CC(work) | | | | |
| CC(free) | | | | |
| Number of non-hydrogen atoms | 4294 | 5346 | 1260 | 1277 |
| macromolecules | 4294 | 5157 | 1217 | 1203 |
| ligands | 0 | 0 | 43 | 51 |

| | | | | |
|---|---|---|---|---|
| solvent | 0 | 189 | 0 | 23 |
| Protein residues | 563 | 687 | 161 | 159 |
| Nucleic acid bases | | | | |
| RMS(bonds) | 0.003 | 0.002 | 0.005 | 0.002 |
| RMS(angles) | 0.6 | 0.56 | 0.77 | 0.63 |
| Ramachandran favored (%) | 95.38 | 98.07 | 98.09 | 98.06 |
| Ramachandran allowed (%) | 4.62 | 1.93 | 1.91 | 1.94 |
| Ramachandran outliers (%) | 0 | 0 | 0 | 0 |
| Rotamer outliers (%) | 1.08 | 0.91 | 0.77 | 0 |
| Clashscore | 4.96 | 2.98 | 2.77 | 1.19 |
| Average B-factor | 71.18 | 29.88 | 51.27 | 42.98 |
| macromolecules | 71.18 | 30.1 | 51.14 | 42.71 |
| ligands | | | 54.81 | 48.84 |
| solvent | | 23.83 | | 44.06 |
| Number of TLS groups | | | | |

**Table S3: Data collection and refinement statistics for GII-Pro R112A mutants in complex with rupintrivir**

| | HOV_R112A_Rupintrivir | GII3_R112A_Rupintrivir | Sydney_R112A_Rupintrivir |
|---|---|---|---|
| PDB ID | 9DA0 | 9DAL | 9D9T |
| Wavelength | | | |
| Resolution range | 46.55 - 2.28 (2.61 - 2.28) | 41.16 - 2.45 (2.58 - 2.45) | 42.59 - 2.4 (2.75 - 2.4) |
| Space group | P 32 2 1 | P 32 2 1 | P 32 2 1 |
| Unit cell | 93.1 93.1 44.66 90 90 120 | 95.05 95.05 46.18 90 90 120 | 98.35 98.35 46.28 90 90 120 |
| Total reflections | 120094 (40176) | 157911 (18573) | 231754 (78563) |
| Unique reflections | 10832 (3558) | 9064 (1274) | 10470 (3455) |
| Multiplicity | 11.1 (11.3) | 17.4 (14.6) | 22.1 (22.7) |
| Completeness (%) | 92.03 (100.00) | 99.96 (100.00) | 90.82 (85.13) |
| Mean I/sigma(I) | 16.76 (3.50) | 22.49 (0.97) | 25.42 (12.06) |
| Wilson B-factor | 41.62 | 71.56 | 44.47 |
| R-merge | 0.09514 (0.7532) | 0.08889 (2.865) | 0.08203 (0.2326) |
| R-meas | 0.09999 (0.7897) | 0.09151 (2.968) | 0.08396 (0.2378) |
| R-pim | 0.03033 (0.2359) | 0.02146 (0.7685) | 0.0177 (0.04935) |
| CC1/2 | 0.995 (0.924) | 1 (0.352) | 0.999 (0.994) |
| CC* | 0.999 (0.98) | 1 (0.722) | 1 (0.999) |

| | | | |
|---|---|---|---|
| Reflections used in refinement | 9589 (3410) | 9060 (1274) | 9383 (2892) |
| Reflections used for R-free | 480 (170) | 913 (128) | 473 (142) |
| R-work | 0.1985 (0.2614) | 0.2237 (0.3549) | 0.2172 (0.2954) |
| R-free | 0.2330 (0.2626) | 0.2484 (0.3700) | 0.2512 (0.3227) |
| CC(work) | | | |
| CC(free) | | | |
| Number of non-hydrogen atoms | 1323 | 1280 | 1294 |
| macromolecules | 1242 | 1237 | 1204 |
| ligands | 43 | 43 | 47 |
| solvent | 38 | 0 | 43 |
| Protein residues | 165 | 165 | 160 |
| Nucleic acid bases | | | |
| RMS(bonds) | 0.002 | 0.017 | 0.003 |
| RMS(angles) | 0.68 | 0.75 | 0.63 |
| Ramachandran favored (%) | 97.52 | 98.14 | 98.08 |
| Ramachandran allowed (%) | 2.48 | 1.86 | 1.92 |
| Ramachandran outliers (%) | 0 | 0 | 0 |
| Rotamer outliers (%) | 0 | 1.52 | 0.78 |
| Clashscore | 1.93 | 2.33 | 0.4 |
| Average B-factor | 46.19 | 80.84 | 42.04 |
| macromolecules | 46.31 | 80.83 | 42.01 |
| ligands | 44.73 | 80.98 | 43.21 |
| solvent | 44.24 | | 41.75 |
| Number of TLS groups | | | |

**Table S4: Crystallization conditions**

| **Crystal** | **Condition** |
|---|---|
| GII.3 Apo | 0.1-0.35M KSCN, 15-25% PEG 3,350, 0.1M Bis-Tris Propane pH 7.5 |
| GII.4 Sydney Apo | 1-8% Tacsimate pH 5.0, 4-16% PEG 3,350 |
| GII.3 - Rupintrivir | 0.3 M Sodium Formate, 0.1 M Sodium Citrate Tribasic:HCl, 22.5 % (v/v) PurePEGs Cocktail |
| GII.3 R112A - Rupintrivir | 0.1 M BICINE pH 9.0, 10% (w/v) PEG 20000; 2%(v/v) 1,4-Dioxane |
| GII.4 Sydney - Rupintrivir | 0.3 M Sodium Iodide, 0.1 M Potassium Nitrate:NaOH, 22.5 % (v/v) PurePEGs Cocktail |
| GII.4 HOV - Rupintrivir | 0.1 M BIS-TRIS pH 5.5, 2.0 M Ammonium sulfate |
| GII.4 HOV R112A - Rupintrivir | 0.2 M Lithium acetate dihydrate, 20% w/v Polyethylene glycol 3,350, pH 7.9 |
| GI.1 - Rupintrivir | 0.2 M Lithium sulfate, 0.1M Tris, 1.0 M Sodium/Potassium tartrate, pH 7.0 |

| | |
|---|---|
| GII.4 R112A - Rupintrivir | 0.02 M Magnesium chloride hexahydrate, 0.1M HEPES pH 7.5, 22 % w/v Poly(acrylic acid sodium salt) 5100, 0.2M potassium iodide |
